# *Vibrio cholerae* motility in aquatic and mucus-mimicking environments

**DOI:** 10.1101/2021.07.06.451398

**Authors:** Marianne Grognot, Anisha Mittal, Mattia Mah’moud, Katja M. Taute

## Abstract

Cholera disease is caused by *Vibrio cholerae* infecting the lining of the small intestine and results in severe diarrhea. *V. cholerae*’s swimming motility is known to play a crucial role in pathogenicity and may aid the bacteria in crossing the intestinal mucus barrier to reach sites of infection, but the exact mechanisms are unknown. The cell can be either pushed or pulled by its single polar flagellum, but there is no consensus on the resulting repertoire of motility behaviors.

We use high-throughput 3D bacterial tracking to observe *V. cholerae* swimming in buffer, in viscous solutions of the synthetic polymer PVP, and in mucin solutions that may mimic the host environment. We perform a statistical characterization of its motility behavior on the basis of large 3D trajectory datasets. We find that *V. cholerae* performs asymmetric run-reverse-flick motility, consisting of a sequence of a forward run, reversal, and a shorter backward run, followed by a turn by approximately 90°, called a flick, preceding the next forward run. Unlike many run-reverse-flick swimmers, *V. cholerae*’s backward runs are much shorter than its forward runs, resulting in an increased effective diffusivity. We also find that the swimming speed is not constant, but subject to frequent decreases. The turning frequency in mucin matches that observed in buffer. Run-reverse-flick motility and speed fluctuations are present in all environments studied, suggesting that these behaviors may also occur in natural aquatic habitats as well as the host environment.

**Importance:** Cholera disease produces vomiting and severe diarrhea and causes approximately 100,000 deaths per year worldwide. The disease is caused by the bacterium *Vibrio cholerae* colonizing the lining of the small intestine. *V. cholerae*’s ability to swim is known to increase its infectivity, but the underlying mechanisms are not known. One possibility is that swimming may aid in crossing the protective mucus barrier that covers the lining of the small intestine. Our work characterizing how *V. cholerae* swims in environments that mimic properties of the host environment may advance the understanding of how motility contributes to infection.

## Introduction

Motility has been recognized as a major virulence factor in *V. cholerae*(1, 2), the causative agent of cholera disease, which produces severe watery diarrhea. The *V. cholerae* population recovered from the so-called “rice-water” stool of infected patients shows both stronger motility and greater infectivity than the same strain grown in the lab(3), and non-motile *V. cholerae* mutants show decreased infectivity(4–6). The flagella that drive motility are a key target of the host immune response: flagellins trigger inflammatory responses from the host(7), and immune infant mice carry antibodies that specifically inhibit flagellum-mediated motility(8). Motility has been suggested to aid the bacteria in crossing the intestinal mucus barrier to access sites of colonization(4, 6, 9).

Despite the impact of motility on pathogenicity, characterizations of *V. cholerae*’s motility behavior have largely been qualitative, and it is commonly described as having the appearance of “shooting stars” in darkfield microscopy(10–12). Its single polar flagellum allows the cell to swim forward or backward, being pushed or pulled by the flagellum, respectively, depending on the direction of flagellar rotation. The shorter backward swimming segments have been reported to be accompanied by random reorientation(2, 13). This description is reminiscent of the well-studied “run-tumble” motility of *Escherichia coli*, where swimming (“running") is driven by counterclockwise (CCW) rotation of the bundled flagella, while intermittent clockwise (CW) rotation by one or more flagella breaks up the bundle and results in short reorientation events called “tumbles”. In line with this, expected motility phenotypes of chemotaxis gene deletion mutants in *V. cholerae* have often been assigned by analogy to *E. coli*(2, 13, 14). More recent work(15, 16) has described *V. cholerae*’s motility as the “run-reverse-flick” behavior reported for other polarly flagellated Vibrio species(17–19), where the transition from pushing to pulling is accompanied by a reversal, while the opposite transition is accompanied by a turn of approximately 90° called a “flick”. These two views seem superficially similar, but differ in their prediction for how chemotaxis, that is, the bacterium’s ability to bias its direction of motion relative to chemical gradients, can be achieved. Neither view has been supported by a full quantitative characterization of *V. cholerae*’s motility behavior.

Such a characterization, however, is crucial to understanding how motility and chemotaxis contribute to pathogenicity. The role of chemotaxis in pathogenicity is still debated(1). Butler and Camilli(2, 13) demonstrated that the infectivity of chemotaxis mutants depends on their motility phenotype, underscoring the need for an accurate, quantitative assessment of such phenotypes. Currently such an assessment is lacking even for wildtype strains.

Here we use high-throughput 3D tracking to demonstrate that *V. cholerae* performs run-reverse-flick motility with variable swimming speed in aquatic environments and quantify the associated behavioral parameters. We show that its motility strategy of asymmetric forward and backward run durations enhances diffusive spreading. To determine whether *V. cholerae* motility behavior differs in environments that mimic physical properties of the host environment, we also observe *V. cholerae* motility in viscous polymer solutions. We show that this behavior is retained in environments that approximate the physical complexity of natural habitats.

## Results

### Vibrio cholera performs run-reverse-flick motility in aquatic environments

We utilize a high-throughput 3D bacterial tracking technique(20) to gather large datasets of 3D trajectories for swimming Vibrio cholerae O395-NT(21) (toxin deletion in classical biotype strain O395) traversing a volume spanning hundreds of µm in each direction, tens of µm from the sample chamber surfaces (Fig 1a). We analyze 23,062 3D trajectories with an average swimming speed greater than 20 µm/s and a duration above 1 s, cumulating in 58,029 s of trajectory time, from three biological replicates. The average swimming speed in motility buffer is 94 µm/s (Fig. 1b, Supplementary Figure 1).

**Figure 1:**
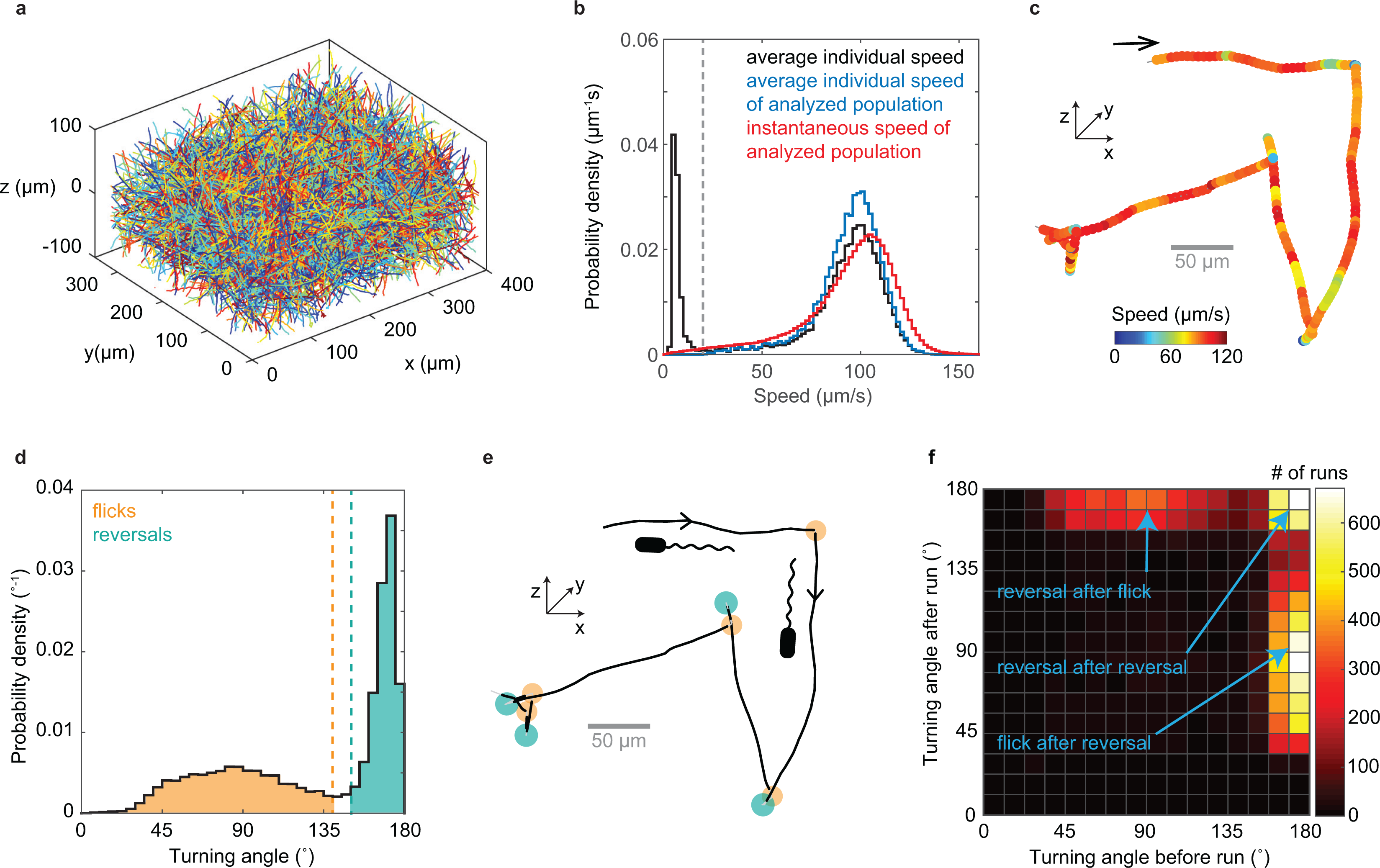
*V. cholerae* 3D motility characterization. a) 3D trajectories obtained from one 100 s-long video recording. b) Probability distribution of average individual swimming speeds of the full population (black, weighted by trajectory duration) and of instantaneous swimming speeds for the analyzed population (red), consisting of trajectories with an average speed larger than 20 µm/s (marked by dashed line) and a minimal duration of 1 s. c) Example trajectory with color reflecting swimming speed. The arrow marks the trajectory start. d) Distribution of turning angles and classification of turn events. Turns by less than 140° are considered flicks, and those by more than 150° are considered reversals. Flick and reversal angles have magnitudes of 88° ± 26° (mean ± SD) and 169° ± 6°, respectively. e) Turn event identification in the trajectory from panel c reveals alternating flicks (orange) and reversals (teal). f) Bivariate histogram of consecutive turning angles observed after versus before the same run. Reversals can be preceded and followed by reversals or flicks, but two flicks never occur in a row.

Visual inspection of individual 3D trajectories reveals approximately straight runs, bordered by turns that alternate in magnitude, indicative of run-reverse-flick motility (Fig. 1c). We employ an automated procedure to detect turning events (Methods). Similar to previous reports for run-reverse-flick motility in *Vibrio alginolyticus*(20) and *Caulobacter crescentus*(22), the turning angle distribution (Fig. 1d) shows two distinct peaks, one narrow peak near 180°, reflecting reversals, and a second broader peak with a center near 90°, which we attribute to flicks.

Flicks are thought to be caused by a buckling instability of the flagellar hook that connects the flagellar filament to the flagellar motor(18). During backward swimming, the flagellum pulls the cell, and the hook is likely stretched out. When the cell switches to forward swimming, the hook is compressed by the pushing flagellum and can buckle under the load if a critical force or torque threshold is reached. The buckling typically occurs within approximately 10 ms after motor reversal(18). This delay between reversal and buckling-induced reorientation is not resolved in our video rate recordings, thus the observed flick encompasses both events. If no buckling occurs, only a reversal is observed. Thus, multiple reversals may occur in sequence. Because flicks only occur during the transition from backward to forward swimming, two flicks cannot occur in a row. To test these predictions, we analyze the magnitudes of consecutive turns (Fig. 1f) and find that reversals and flicks typically alternate. A reversal can also be followed by another reversal, but we do not observe consecutive flicks. The observed pattern of turning angle magnitudes is thus consistent with run-reverse-flick motility.

*V. cholerae*’s flagellar motor is driven by a sodium motive force(23). Decreasing the sodium motive force by decreasing the sodium concentration, [Na+], at constant ionic strength, results in a lower swimming speed (Supplementary Figure 1f). The concomitant decrease in the occurrence of flicks (Fig. 2) is consistent with the motor torque driving the underlying buckling transition(18). We thus conclude that *Vibrio cholerae* performs run-reverse-flick motility.

**Figure 2:**
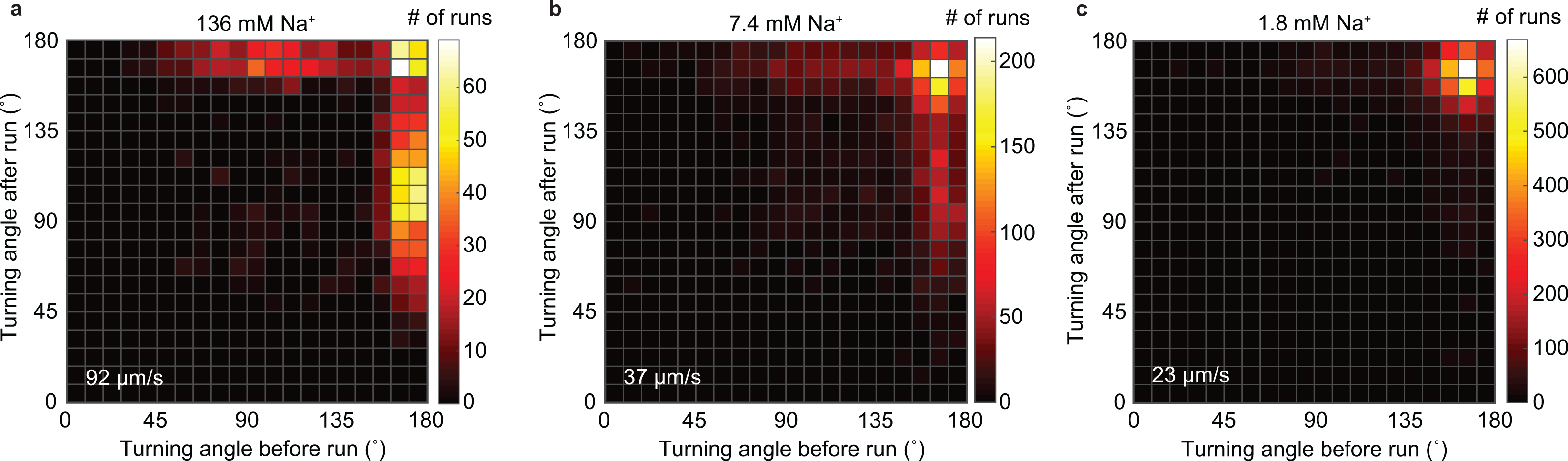
*V. cholerae* flicking probability depends on the sodium motive force. Bivariate histograms of consecutive turning angles for sodium concentrations of 136 mM (a), 7.4 mM (b) and 1.8 mM (c), with respective average swimming speeds of 92, 37, and 23 µm/s. The sodium concentration for Fig. 1f is 181 mM and the average swimming speed 94 µm/s.

### Forward runs are longer than backward runs

Identifying turn events as flicks and reversals by their magnitude enables us to assign a bacterial orientation to runs without the need to visualize the flagellum. Of the 18,533 complete runs in our dataset, we can identify 4,581 as forward and 8,180 as backward (Methods). Forward runs show right-handed trajectory curvature when swimming along the bottom surface of the sample chamber, consistent with a left-handed flagellum pushing the cell by CCW rotation(24) (Supplementary Figure 2). We obtain similar average swimming speeds for forward and backward runs of 90 µm/s and 86 µm/s, respectively. Run duration distributions for both directions show a peak at approximately 0.08 s, followed by an approximately exponential decay which is rapid for backward runs and slow for forward runs (Fig. 3b). Peaked run duration distributions have also been found for other polarly flagellated bacteria, *Vibrio alginolyticus*(17) and *Caulobacter crescentus*(25). On average, forward runs are approximately 3.6 times longer than backward runs. While we observe no correlation in the duration of consecutive forward and backward runs (Fig. 3c), the differences in the duration of forward runs and their subsequent backward runs are exponentially distributed (Fig. 3d), with different time scales dependent on which run type is longer. One key feature of run-reverse-flick motility is that backward runs roughly retrace the preceding forward run. Given similar forward and backward run speeds, the net displacement caused by one such pair of runs is thus determined by their difference in durations. Exponentially distributed run duration differences have also been observed for *V. alginolyticus*(17) and thus may represent a common feature of species performing run-reverse-flick motility.

**Figure 3:**
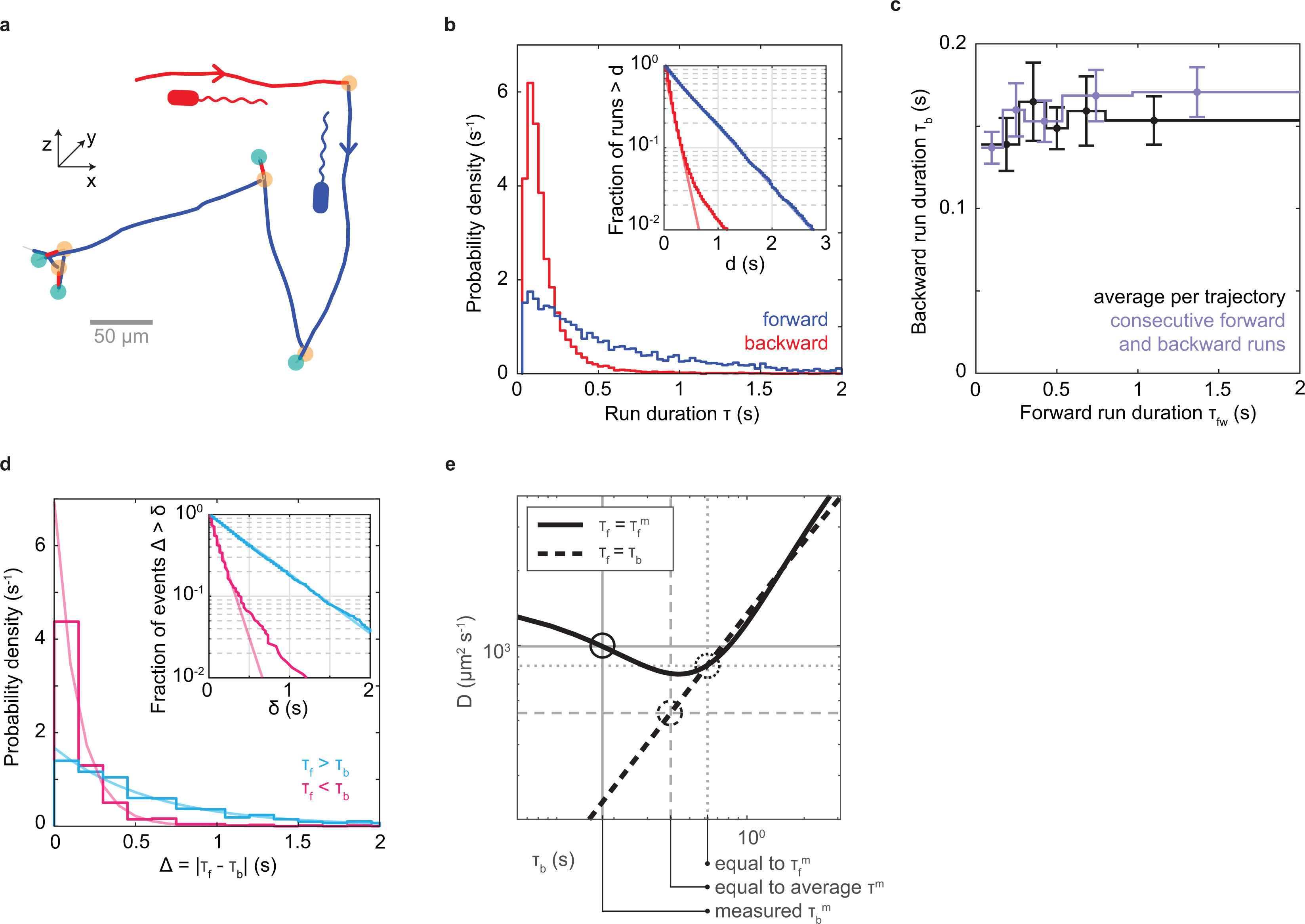
Run characterization. a) Example trajectory from Fig. 1c with runs identified as forward (blue) or backward (red) based on the identity of the bordering turning events (orange: flicks, teal: reversals). b) Run duration distributions. Average durations of backward (red) and forward (blue) runs are 0.174 ± 0.002 s (mean ± SE) and 0.62 ± 0.01 s, respectively. Inset: Fraction of runs that are longer than a threshold, d, as a function of d. Line fits in log-linear space to the ranges of 0.1-1.5 s and 0.067-0.4 s yield exponential decay time scales of 0.60 s and 0.14 s for forward and backward runs, respectively. The tail of the backward run duration distribution likely represents misidentified forward runs. c) Relationship between backward and forward run duration. Purple: Average duration of backward runs as a function of the preceding forward run’s duration; black: average duration of backward runs as a function of average duration of forward runs for trajectories containing at least 4 runs of known orientation. d) Distribution of the absolute differences in duration between a forward run and the subsequent backward run, shown in cyan (magenta) when the forward run is longer (shorter) than the backward. Inset: Fraction of events that are longer than a threshold, x, as a function of x. Partially transparent lines indicate exponential decay fits. For positive differences (cyan), a maximum likelihood fit of an exponential distribution yields an exponential decay time of 0.60 s. For negative differences (magenta), the slope of a line fit in semi-log space in the range of 0-0.8 s yields a decay time of 0.14 s. We attribute the tail of the distribution for negative differences to misidentified forward runs, as in panel b. e) Predicted effective diffusion coefficient for run-reverse-flick motility as a function of the average backward run duration, τ_b_, based on results of Taktikos et al.(26) (see Supplementary Note 1 for details). The black solid line indicates a fixed forward run duration equal to the measured value, τ_f_ = τ_f_^m^ = 0.62 s, and the black dashed line indicates equal forward and backward run durations. Grey dashed lines mark a scenario where both forward and backward run durations are equal to the average measured run duration. Gray dotted lines mark the scenario where both equal the measured forward run duration. Grey solid lines indicate the measured scenario of τ_f_^m^ = 0.62 s and τ_b_^m^ = 0.174 s.

Run durations are expected to affect random spreading of cells(26) (Fig. 3e). The run-reverse-flick swimmers *V. alginolyticus* and *C. crescentus* show similar durations of forward and backward runs(17, 22). Theoretical work allows a prediction of the effective diffusion coefficient characterizing random spreading at long time scales(26). *V. cholerae*’s average run duration is similar to that of *V. alginolyticus*, but its asymmetry in forward and backward run durations is expected to yield an 85% increase in the diffusion coefficient, compared to a symmetric scenario with the same average run duration (Fig. 3e, Supplementary Note 1). The diffusion coefficient is also enhanced by 19% compared to a scenario where both run durations equal the longer, forward run duration. We thus conclude that *V. cholerae*’s asymmetric run durations enhance random spreading of cells.

### Run speed modulation

We observe that the swimming speed during run segments is not constant, but subject to substantial, temporary decreases that are readily apparent by visual inspection of trajectories (Fig. 4a). During these events, the bacterium maintains its previous swimming direction (Fig. 4b) but follows it at a decreased speed. We employ an automated detection procedure that identifies these deceleration events as decreases in speed below a threshold that last for at least two consecutive frames. The speed threshold is set at a fixed fraction β of a run-specific baseline value (Methods), and we only consider runs with a minimum duration of 10 frames (0.33 s) to ensure that a baseline speed value can be confidently assigned. Although we consider runs of either orientation in the analysis, the runs that meet the duration threshold are likely primarily forward runs. At the selected value of β = 0.75, we obtain a deceleration event frequency of 0.53 Hz during runs (Methods). While their frequency is similar to the turning frequency, we detect no strong correlation between estimates of the deceleration frequency and the turning frequency obtained for individual bacteria (Supplementary Figure 4f). The durations of the deceleration events are exponentially distributed (Fig. 4c), with an average duration of 0.12 s, suggesting that escape from these events may be a random event with a fixed rate of occurrence. The intervals between decelerations, however, show a peaked distribution with a maximum at 0.12-0.15 s, followed by an exponential tail (Fig. 4d).

**Figure 4:**
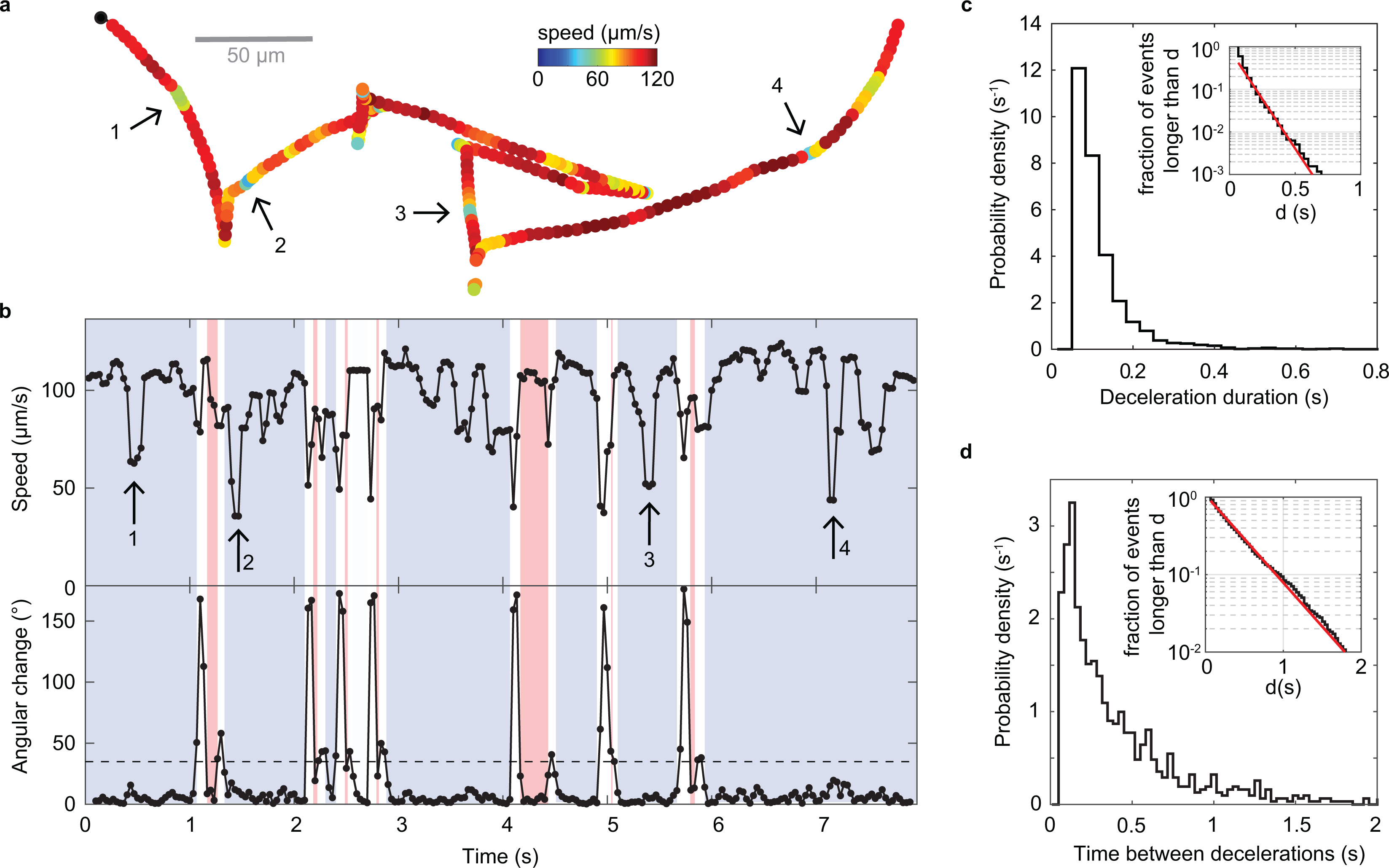
Deceleration events. a) Example trajectory with visually apparent segments of decreased speed marked by arrows. b) Time series of swimming speed (top) and angular change in swimming direction between consecutive frames (bottom) for the trajectory shown in panel a. Blue segments represent forward swimming, red segments backward swimming. c) Distribution of durations of deceleration events. The average duration is 0.12 s. Inset: Fraction of events longer than a threshold, d, as a function of d. The slope of a linear fit (red) in in semi-log space yields an exponential decay time of 0.094 s. d) Distribution of time between two consecutive decelerations. The average is 0.41 s. Inset: Fraction of events longer than a threshold, d, as a function of d. The slope of a linear fit (red) in the range of 0.16-2 s in semi-log space yields an exponential decay time constant of 0.38 s.

To determine whether these drops in speed represent discrete events or a continuous variation in speed, we examine the dependence of their properties on the event detection threshold. Their frequency, duration, and relative speed distributions vary continuously with the threshold parameter, β (Supplementary Figure 4c-e, f), suggesting that they may derive from a continuous variation in speed. The ratio of variances of instantaneous and trajectory-averaged speeds of the analyzed population provides an upper limit on the contribution of inter-individual, as opposed to temporal, variability at the population level. Based on the standard deviations of 24 µm/s and 17 µm/s observed for their respective distributions, we conclude that at least half the variability in speed observed at the population level derives from such temporal, intra-individual variability.

Errors in the bacterial position determination can produce fluctuations in the measured velocity if the magnitude of the errors is comparable to the true displacement between neighboring frames. The errors on the measured speed are thus larger for slower swimming bacteria. To rule out measurement errors as the source of the observed speed fluctuations, we acquire trajectories for another run-reverse-flick swimmer with a lower swimming speed, *Caulobacter crescentus*, under the same measurement conditions. We find that the relative variation in run speed is larger for *V. cholerae* than for *C. cresce*ntus (Supplementary Figure 4g) and thus rule out measurement errors as the cause of the speed variations observed in *V. cholerae*. We also confirm that deceleration events are not caused by our trajectory filtering method (Supplementary Figure 4a, b).

### Run-reverse-flick motility in mucin solutions

Run-reverse-flick motility has been viewed as an adaptation to marine habitats(27), raising the question of whether this behavior is preserved in environments as physically complex as the host environment where the bacteria have to cross a viscous mucus barrier that protects the intestinal epithelium to reach sites of infection.

The primary component of mucus are mucins, large glycoproteins which form a hydrogel with an estimated physiological concentration range of 1-5%(28). We track *V. cholerae* swimming in solutions of 1.2% mucin purified from human saliva. Run-reverse-flick motility is still readily apparent from the trajectories (Fig. 5a). We analyze 2,327 3D trajectories of at least 1 s duration and 20 µm/s average speed. The swimming speed has decreased to 57 µm/s (Fig. 5b). The bivariate distribution of turn angle magnitudes before and after a run indicates alternating flicks and reversals, or sequences of reversals, consistent with run-reverse-flick motility (Fig. 5d). The magnitude of the flicks has increased from 88°± 26° (mean ± SD, 10,946 angles) observed in buffer to 97°± 33° (mean ± SD, 1,046 angles). As our flick angle measurements combine a reversal and a reorientation occurring in rapid succession, the increased flick angle indicates a decreased reorientation, consistent with increased drag. The fraction of turn events identified as reversals is very similar in the absence and presence of mucin (Fig. 5e). While the fraction of turn events identified as flicks is slightly lower in mucin compared to in buffer, we attribute this difference the increased difficulty of identifying flicks due to their increased magnitude. We thus conclude that the flick probability is likely very similar in the presence of mucin than in its absence. The observed event rates are compatible with flick probabilities in the range of 88-93% in the absence and 81-94% in the presence of mucus, depending on whether the unidentified turn events are dominated by flicks or reversals.

**Figure 5:**
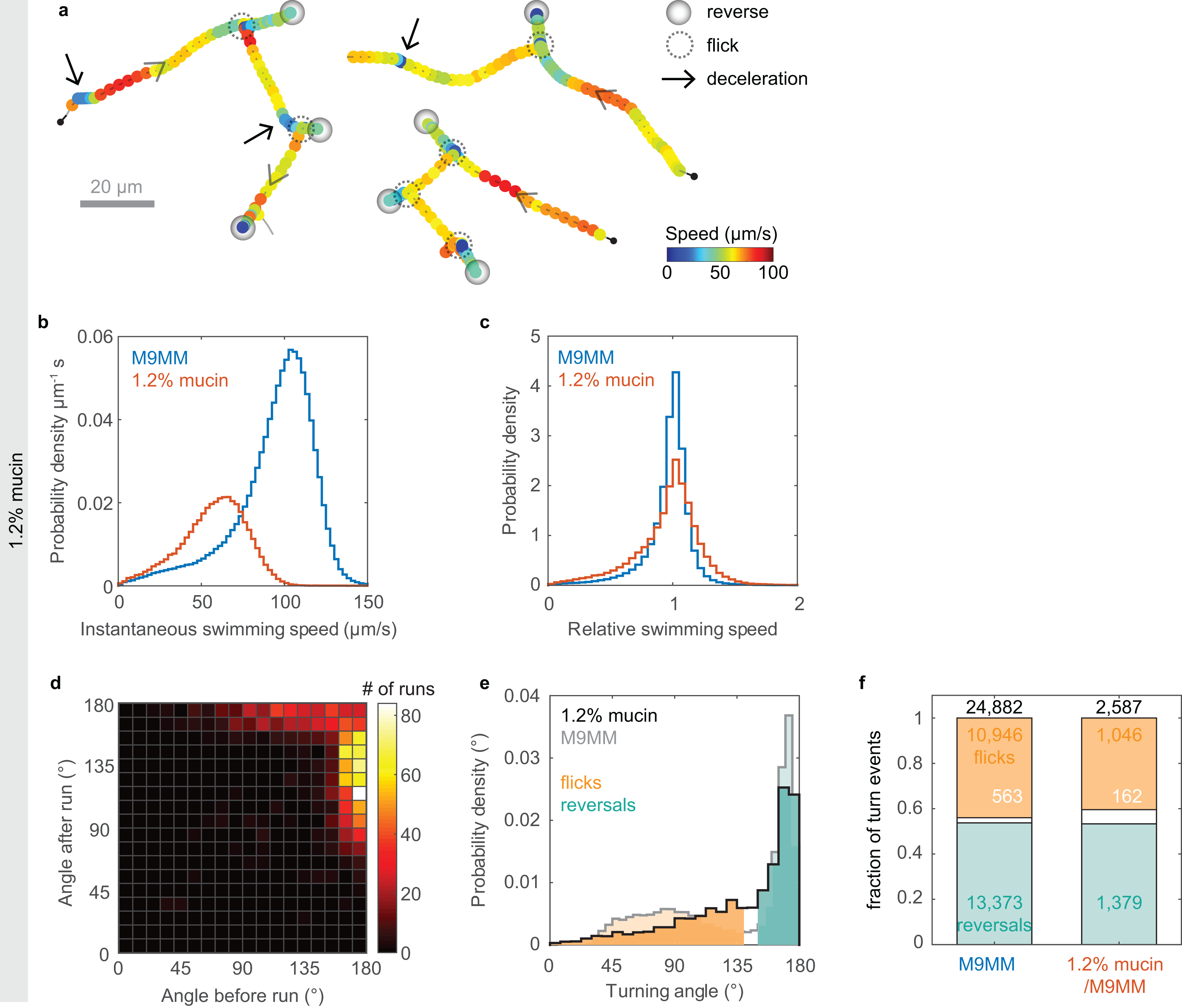
Run-reverse-flick motility in solutions of 1.2% mucin in M9MM. a) Three example trajectories in 1.2% mucin, with marked reverse, flick, and deceleration events. b) Distribution of instantaneous swimming speeds observed in the presence (red) and absence (blue) of mucin. The average speeds are 57 µm/s in mucin and 94 µm/s in M9MM. c) Distribution of relative swimming speeds in the presence (red) and absence (blue) of mucin. The relative speed is the instantaneous swimming speed divided by the individual’s median swimming speed. d) Bivariate histogram of consecutive turning angles in 1.2% mucin also displays alternating flicks and reversals as well as consecutive reversals, consistent with run-reverse-flick motility. e) Distribution of turning angles and classification of turn events for trajectories in M9MM (grey, reproduced from Fig. 1d) and in mucin (black). The average flick angle is 88° ± 26° (mean ± SD) in M9MM and 97°± 33° in 1.2% mucin/M9MM. The average reversal angle is 169° ± 6° in M9MM and 168 ± 7° in mucin. f) Fraction of turn events classified as flicks (orange), reversals (teal), or unidentified (gray) in the absence and presence of 1.2% mucin. Turns of an angle up to 140° are considered flicks, those above 150° reversals, and those in between 140° and 150° are considered unidentified. Absolute numbers of events for each category are given.

The observed turning frequency of 0.49 Hz in mucin matches that observed in buffer. The backward and forward run durations of (0.17±0.01) s (mean ± SE) and (0.71±0.04) s, respectively, are also close to the values of (0.174 ± 0.002) s and (0.62±0.01) s observed in buffer. We also still observe temporary decreases in swimming speed during runs (Fig. 5a) but cannot rule out that inhomogeneities in the mucin solution contribute to them. The relative variability in swimming speeds is slightly increased in the presence of mucin, with a coefficient of variation, defined as the ratio of standard deviation and mean, of relative swimming speeds of 0.29 in mucin and 0.25 in buffer (Fig. 5c).

### Run-reverse-flick motility is preserved in dense polymer solutions

The gut environment likely also contains regions that are denser than the dilute mucin solutions used here. To determine whether *V. cholerae* motility behavior differs qualitatively in denser environments, we observe *V. cholerae* motility in solutions of the synthetic, large molecular-weight polymer PVP K90 (Figure 6). Run-reverse-flick motility is still apparent even at macroscopic viscosities more than 50 times that of water where the swimming speed has dropped to 11 µm/s (Fig. 6a-c). With increasing polymer concentration, the associated changes in refractive index cause increasing localization errors. Because these errors can cause ambiguities in trajectory interpretation, we refrain from performing a quantitative turning analysis at this point. Variations in swimming speed during runs are still visually apparent (Fig. 6a-c) and occur at a similar relative amplitude as in buffer (Fig. 6e).

**Figure 6:**
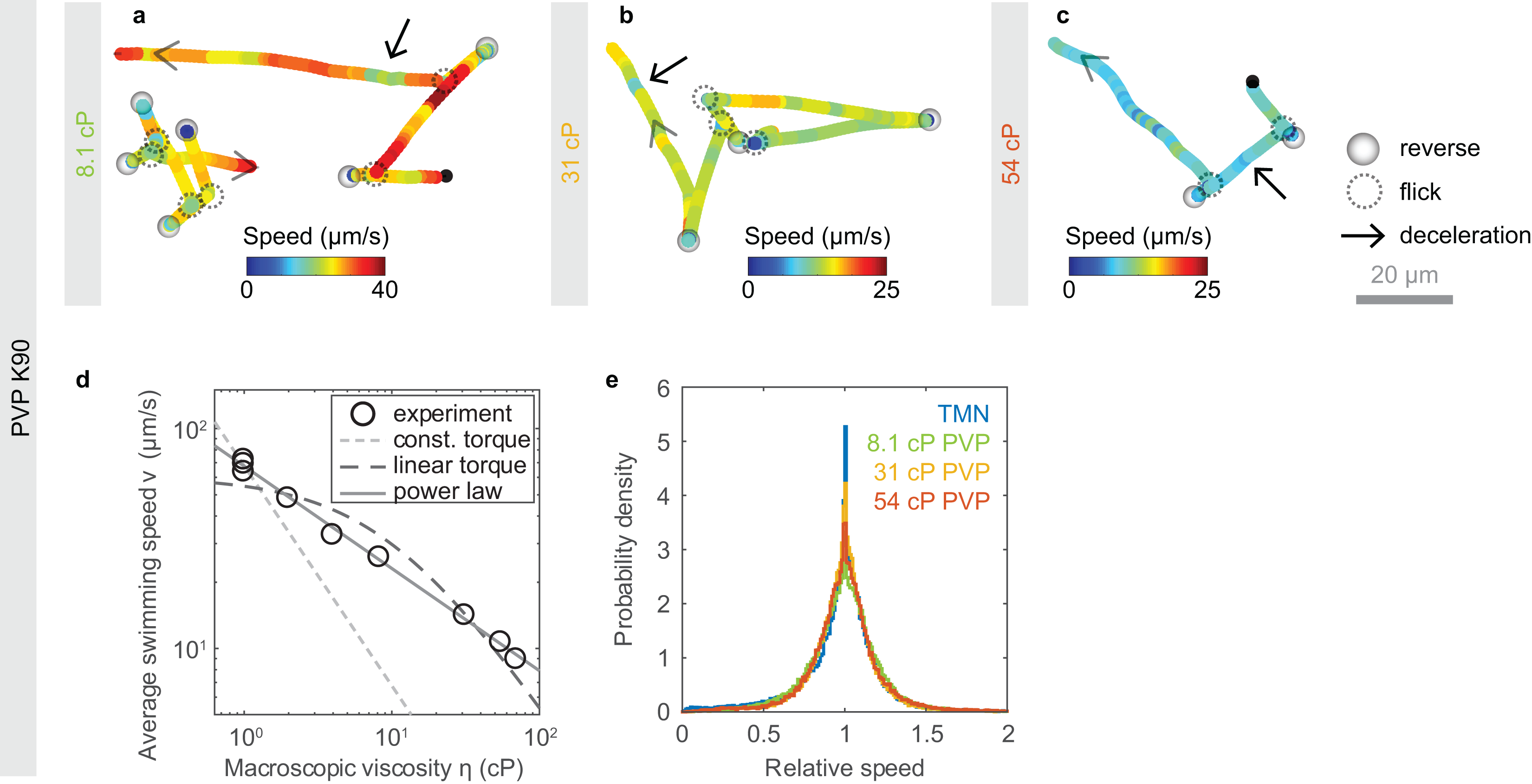
Run-reverse-flick motility in PVP K90 solutions. a-c) Example trajectories showing run-reverse-flick motility and variations in swimming speed at concentrations of 2.2%, 4.5%, and 6% PVP K90 in TMN, respectively. d) Average swimming speed as a function of macroscopic PVP K90 viscosity. The dashed and dotted lines represent two different motor torque models under the assumption of Newtonian fluid behavior. The light gray, dotted line indicates a η^-1^ dependence corresponding to a constant motor torque. The darker dashed line is a fit of the dependence v = a / ( η + b) that is expected for a linear torque-speed relationship(40). Parameters a = 588 µm cP s^-1^ and b = 9.7 cP yield the best error-weighted fit when a fixed relative measurement error in swimming speed is assumed. The gray solid line represents a power law fit with a dependence η^-0.46^. e) Distribution of relative swimming speeds for different PVP concentrations, reflecting constant variability of swimming speeds. The relative speed is the instantaneous swimming speed divided by the individual’s median swimming speed.

In a Newtonian liquid with viscosity η, the swimming speed would be expected to decay as 1/ η if the torque of the flagellar motor remains constant. The average swimming speed of *V. cholerae*, however, decreases more slowly with the macroscopic viscosity of PVP K90 solutions (Fig. 6d). The flagellar motors of *E. coli*(29) and *V. alginolyticus*(*30*) have been shown to exhibit a complex torque-speed relationship that can be approximated by a constant torque regime at low speeds and high loads and a linear decrease in torque as a function of rotation speed beyond a rotation speed threshold called the knee frequency. A linear torque-speed relationship, in combination with Newtonian hydrodynamics, however, does not provide a good fit to the data, either (Fig. 6d). In both cases, the measured swimming speeds at high macroscopic viscosity exceed the theoretical expectation. Empirically, a power law fit with an exponent of 0.46 provides a good approximation to the data.

We conclude that run-reverse-flick motility as well as speed variations are present in solutions of both synthetic and natural polymers, and thus may also occur in the host environment.

### Strain dependence of observations

While we performed our experiments with strain O395-NT(21), a mutant of classical strain O395 that carries a deletion of both subunits of the cholera toxin (ctxAB), visual inspection of trajectories obtained for wildtype O395 reveals no differences in behavior (Supplementary Figure 3). Both run-reverse-flick motility and speed modulation during runs are present in both strains. Automated trajectory analysis reveals similar turning frequencies as well as very similar average backward and forward run durations for strains O395-NT and wildtype O395 (Table 1).

**Table 1:** Comparison of motility parameters between strains O395-NT and wildtype O395 in M9MM. Number of events on which the value is based are shown in brackets.

| Strain | Analyzed trajectory time | Average swimming speed | Turning frequency | Average forward run duration | Average backward run duration |
| --- | --- | --- | --- | --- | --- |
| O395-NT | 58,029 s<br>(23,062) | 94 $\mu\text{m/s}$ | 0.49 Hz<br>(28,703) | $0.62 \pm 0.01$ s<br>(4,581) | $0.174 \pm 0.002$ s<br>(8,180) |
| O395 (wt) | 19,113 s<br>(8,017) | 88.5 $\mu\text{m/s}$ | 0.47 Hz<br>(8,978) | $0.58 \pm 0.02$ s<br>(1,240) | $0.18 \pm 0.01$ s<br>(2,283) |

The observed turning frequency of 0.49 Hz aligns well with the value of approximately 0.6 Hz previously reported(31) for O395-N1 (ΔctxA in O395(21)), but is much higher than the value of 0.14 Hz previously reported for the El Tor strain C6709-1(13). Together with the finding that smooth-swimming mutants possess greater infectivity(13), this discrepancy raises the intriguing question whether selection on turning frequency may have contributed to the displacement of classical strains by El Tor strains in the current *V. cholerae* pandemic.

## Discussion

During its life cycle, *V. cholerae* encounters a wide range of different environments. Outside the host, it is found in freshwater and in brackish waters both as individual, planktonic swimmers and as biofilms that grow, for instance, on the surface of phytoplankton, zooplankton and other chitinous particles(32). Inside the host, the bacterium encounters the complex 3D environments of the digestive system with a wide range of viscosities, porosities, and inhomogeneities. While our observations for *V. cholerae* swimming in buffer likely translate to planktonic cells in natural aquatic environments such as brackish waters, the host environment is not characterized well enough to determine with certainty how well the polymer solutions used in our experiments approximate it. *V. cholerae* colonizes the small intestine, which is lined by a loose, non-attached mucus layer(33, 34). For the colon, which additionally possesses a thick attached mucus layer with an estimated mucin concentration of approximately 6%(35), the concentration of the non-attached, loose layer has been estimated to be approximately four times lower than that of the attached layer(36), so likely similar to the mucin concentration of 1.2% used in our experiments. We thus expect that our mucin solutions may approximate the concentration of the loose mucus layer of the small intestine, though properties of the salivary mucin MUC5B used here may differ from those of intestinal mucins. While detailed rheological data for the MUC5B concentrations used here is not available, literature values on other mucins and more dilute MUC5B solutions suggest a macroscopic viscosity in the range of 2-15 cP (Supplementary Note 2).

In contrast to our finding of a very similar turning frequency and flick probability in buffer and in 1.2 % purified salivary mucin solutions, a recent study(19) reports a decreased turning frequency and decreased flick probability for *V. cholerae* in solutions of 2% porcine gastric mucin, prepared from commercially available, rehydrated dried porcine gastric mucin. Given *V. cholerae*‘s chemotactic attraction to mucus(37), the reported decrease in turning frequency might reflect a transient chemotactic response to the mucin solution. Our experimental approach limits such transient effects by incubating the bacteria in the motility medium with or without mucus for more than 30 min to allow for adaptation to the new environment. In addition, reports that rehydrated dried mucus does not recapitulate characteristic rheological properties of purified mucus(38) suggests that the properties of rehydrated and purified mucin solutions may not be comparable(39). The flick probability is expected to depend on the torque exerted by the flagellar motor and thus drops with decreasing sodium motive force which coincides with a drop in speed(18) (Fig. 2). While, at increasing viscosity, the swimming speed also decreases, the motor torque is expected to remain constant or increase(30). A decreased flick probability is therefore physically not expected at higher viscosity. The recently reported drop in flick probability in mucin(19) might result from the increased difficulty in disambiguating flicks from reversals because of the larger flick angles in mucin.

Work on *E. coli* swimming in PVP K90 solutions indicates that, due to strong shear thinning, the resulting bacterial motility behavior may depend only on macroscopic viscosities, not detailed rheological properties, with the cell body experiencing the solution viscosity and the flagellum the solvent viscosity(40) (Supplementary Note 2). Shear thinning or other non-Newtonian effects may contribute to the slow decrease in swimming speed with macroscopic viscosity we observe in PVP solutions. Additional contributions may arise, however, from the ability of the flagellar motor to remodel itself by increasing the number of torque-generating stators in response to load increases such that different loads result in different torque-speed curves(41, 42). Mucus and mucin also exhibit strong shear thinning(39, 43). The relative decrease in swimming speed we observe in our mucin solutions is similar to that observed in PVP solutions with a macroscopic viscosity of approximately 3 cP (Fig. 6d, Supplementary Note 2). The range of PVP viscosities we cover aligns with the 1 – 30 cP range of macroscopic viscosities determined for mucus from the small intestine(44). Elastic effects are not expected to play a role for either PVP or mucin at the concentrations studied here(40, 45). More generally, our finding that the motility behavior is qualitatively similar in solutions of the natural glycoprotein polymer mucin and over a range of concentrations of the synthetic polymer PVP suggests that our observations may translate to a broader range of polymer solutions.

The swimming speed variations we observe might be related to pauses in swimming which have been reported in *Pseudomonas putida*(46), *Pseudomonas aeruginosa*(*47*), and *Azospirillum brasilense*(48), or to the pauses in flagellar rotation that have been reported in *E. coli*(49), *Rhodobacter sphaeroides*(50), and *P. aeruginosa*(51). For *E. coli*, a correlation between the turning and flagellar pausing frequencies was observed both between mutant strains and between individuals of the same strain(52). We detect no correlation between individual turning and deceleration frequency for *V. cholerae* (Supplementary Figure 4f), but the typically small number of events per trajectory in our data is insufficient to rule out a correlation. We also cannot confidently determine whether deceleration events appear only in forward runs or in both run directions as most backward runs are too short to assign a reliable baseline speed against which variations can be detected. We note that, during the events we observe, swimming never appears to stop, but only decreases in speed, indicating that the flagellum continues to rotate but at a lower speed. In *E. coli*, the speed of flagellar rotation is varied in response to the load on the flagellar motor by changes in the number of torque-producing stators that drive motor rotation(41, 42). Active modulation of flagellar rotation speeds by the chemotaxis signaling system has been reported for proton-driven, unidirectional, peritrichous flagella in *V. alginolyticus*(53), which conditionally expresses these lateral flagella in addition to its polar flagellum, as well as in *Rhizobium meliloti*(54). Transient changes in swimming speed in response to changes in oxygen availability have been reported for *A. brasilense*(55) whose motility is driven by a single, polar, bidirectional flagellum. Tethering experiments may be able to more closely determine the nature and determinants of the modulation in flagellar rotation speed that underlie the swimming speed variation we observe in *V. cholerae*.

*V. cholerae*’s asymmetry in run durations increases its ability to spread randomly (Fig. 3e). In addition, backward swimming segments typically display more trajectory curvature in the vicinity of surfaces than forward swimming segments(56–58) (Supplementary Figure 2). Thus, a bias for forward swimming may further increase dispersal in the vicinity of surfaces compared to a symmetric scenario. While the potential adaptive value of the asymmetric run duration scheme has yet to be evaluated, one possibility is that strong random dispersal is selected for at some point in the *V. cholerae* life cycle. Interestingly, previous work has consistently found that smooth-swimming mutants which suppress turning outcompete the wildtype during infection(4, 13, 59, 60). Such mutants are also expected to display a strongly enhanced effective diffusion coefficient. Another possibility is that asymmetric run durations might have a favorable effect on chemotactic ability, though Altindal et al.(27) have argued that symmetric run durations maximize the chemotactic drift velocity in *V. alginolyticus*.

Run-reverse-flick motility with very short backward swimming segments has also been observed in *Shewanella putrefaciens*(61) and bears some similarities to *E. coli*’s run-tumble motility, where CW rotation does not produce locomotion, but reorientation. These two mechanisms, however, differ in how the degree of reorientation can be controlled. In the run-tumble-like scenario, the magnitude of the reorientation can be increased by prolonging backward run segments, or CW rotation intervals, akin to the relationship between tumble duration and tumble angle that has been observed in *E. coli*(62). In run-reverse-flick motility, by contrast, the magnitude of flick angles is fixed for each individual by the hydrodynamic drag

acting on it(20) and is thus expected to be independent of the duration of CW rotation intervals. The average amount of reorientation, however, also depends on the flicking probability determined by the flagellar motor torque, which can vary with environmental conditions such as salinity (Fig. 2) or nutrient concentrations(63). These differences are likely to affect both random motility and chemotaxis. For instance, the chemotactic precision of the run-reverse-flick bacterium *V. alginolyticus* has been shown to be enhanced by the chemoattractant-dependent modulation of the flicking probability(63) via the swimming speed.

Future work should address *V. cholerae*’s chemotactic mechanism. The well-studied chemotactic strategy of *E. coli* consists of extending average CCW rotation intervals and shortening average CW rotation intervals in response to favorable chemotactic sensory input, thus extending runs up the gradient. The CW bias, that is the fraction of time spent on CW rotation, thus serves as a convenient proxy for the turning frequency which guides chemotaxis. Other species displaying run-reverse-flick motility, *V. alginolyticus*(64) and *C. crescentus*(22), as well as the run-reverse swimmer *P. aeruginosa*(47), instead respond to favorable signals by extending the current flagellar rotation interval regardless of direction, and perform chemotaxis by modulating the turning frequency without substantial change in the bias. While *V. cholerae*’s swimming pattern is closer to the latter category, its short CW rotation intervals are reminiscent of *E. coli*’s short tumbles.

Butler and Camilli(13) termed smooth-swimming chemotaxis mutants CCW-biased and frequently turning mutants CW-biased, based on the well-characterized motility phenotypes of the homologous mutants in *E. coli.*, and attributed their differences in infectivity to the bias in flagellar rotation direction. The bias was, however, not explicitly characterized. Given that bias and turning frequency have been shown to be decoupled in other run-reverse-flick-performing bacteria, and that bias and turning frequency are expected to have distinct effects on motility and chemotaxis(26), future work should address which of these factors drive the observed differences in infectivity. Insights might be gained, in particular, from infectivity assays on smooth-swimming, CW-rotating mutants. We envision that quantitative characterization of motility behaviors of mutants with infectivity phenotypes may present a key tool towards a mechanistic understanding of how motility and chemotaxis behaviors contribute to *V. cholerae* pathogenicity.

## Methods

### Bacterial culturing

Overnight cultures were inoculated from individual *V.* cholerae (O395-NT(21) or O395) colonies, grown on 1.5% agar LB5 plates streaked from glycerol stock, and grown to saturation in 2 ml LB5 (see Table 2 for media compositions) at 30°C, 250 rpm. All media were complemented with 100 µg/ml kanamycin for the O395-NT strain. Day cultures were inoculated at a dilution of 1:200 (v/v) in M9GM and grown at 30°C to an optical density (OD) between 0.350 and 0.400 at 600 nm, unless specified otherwise. While LB5 and TG have previously been used in *V. cholerae* motility studies(5, 13, 23, 65–68), M9 minimal medium with pyruvate (M9GM) yielded higher swimming speeds and lower nonmotile fractions, similar to another recent study(69). We found that an OD range of 0.35-0.4 yielded average swimming speeds above 90 µm/s and a typical motile fraction above 85%.

**Table 2:** Composition of growth and motility media.

|  |  |
| --- | --- |
| LB5 | 1% Bacto Tryptone<br>0.5% Bacto Yeast extract<br>0.5% NaCl<br>pH 7.0 |
| LB10 | 1% Bacto Tryptone<br>0.5% Bacto Yeast extract<br>1% NaCl<br>pH 7.0 |
| TG | 1% Bacto Tryptone<br>0.5% NaCl<br>0.5% (w/v) Glycerol<br>pH 7.1 |
| M9GM | M9 salts (from 5x stock, Sigma)<br>47.7 mM $\text{Na}_2\text{HPO}_4$<br>22 mM $\text{KH}_2\text{PO}_4$<br>8.55 mM NaCl<br>9.35 mM $\text{NH}_4\text{Cl}$<br>0.45 % (77 mM) NaCl<br>0.4 % pyruvate<br>2 mM $\text{MgSO}_4$<br>1 mM $\text{CaCl}_2$<br>pH 7.0 |
| M9MM | M9 salts (from 5x stock, Sigma)<br>47.7 mM $\text{Na}_2\text{HPO}_4$<br>22 mM $\text{KH}_2\text{PO}_4$<br>8.55 mM NaCl<br>9.35 mM $\text{NH}_4\text{Cl}$<br>77 mM NaCl<br>5 mM glucose<br>2 mM $\text{MgSO}_4$<br>1 mM $\text{CaCl}_2$<br>pH 7.0 |
| NoNa-M9MM | 47.7 mM $\text{K}_2\text{HPO}_4$<br>22 mM $\text{KH}_2\text{PO}_4$<br>8.55 mM KCl<br>9.35 mM $\text{NH}_4\text{Cl}$<br>5 mM glucose<br>2 mM $\text{MgSO}_4$<br>1 mM $\text{CaCl}_2$<br>pH 7.0 |
| TMN | 50 mM Tris-HCl<br>300 mM NaCl<br>5 mM $\text{MgCl}_2$<br>5 mM glucose<br>pH 7.5 |
| PYE | 0.2% Bacto Peptone |
|  | 0.1% Bacto Yeast extract<br>1 mM MgSO<sub>4</sub><br>0.5 mM CaCl<sub>2</sub><br>pH 7.0 |

### Sample preparation

Sample chambers with a height of approximately 300 µm were created by using two strips consisting of 3 layers of parafilm as spacers between a microscopy slide and a #1 coverslip. The chamber was then heated on a hot plate and pressed to seal. For samples with mucin, only 2 layers of parafilm were used to reduce sample volumes. Cells were diluted by 1:100 in motility medium M9MM (or fresh growth medium during protocol optimization tests displayed in SI Figure 1a,c), incubated at room temperature for 45 minutes, unless specified otherwise, to allow adaptation to the motility medium, and then flowed into the chamber. Cell solutions are only pipetted with cut pipet tips to avoid shear damage to the flagella. For samples with polymer solutions, cells were first pelleted by centrifugation at 2,000 rcf for 8 minutes, resuspended in the relevant motility medium, diluted 1:200 in the polymer solution, incubated at room temperature for 40-60 minutes, and flowed into the chamber. The ends of the filled chamber were then sealed with molten valap (a mixture of vaseline, lanolin, and paraffin) and immediately brought to the microscope for data acquisition. For motility in M9MM, three to five such samples were prepared and inspected within a period of 10 min for each of three biological replicates for strain O395-NT and one for wildtype O395. For PVP polymer solution experiment, one sample was prepared and inspected per concentration studied, all within a period of 30 minutes. For growth condition tests and mucin solution experiment, two to three samples were prepared and inspected for one biological replicate.

### Data acquisition

Phase contrast microscopy recordings were obtained at room temperature (∼22°C) on a Nikon Ti-E inverted microscope using an sCMOS camera (PCO Edge 4.2, pixel size 6.5 µm) and a 40x objective lens (Nikon CFI SPlan Fluor ELWD 40x ADM Ph2, correction collar set to 1.2 mm to induce spherical aberrations(20)) focused approximately 130 µm above the chamber’s internal bottom surface. The illumination was adjusted to yield approximately 20,000 counts per pixel. Recordings were saved as 16-bit tiff files. For each sample in M9MM, one recording with a duration of 2-2.2 min and a frame rate of 30 fps was obtained immediately after placing the sample on the microscope. The three replicate experiments for O395-NT generated a cumulative 23 min of video recordings. For each PVP sample, one recording was obtained at 30 fps or 15 fps for concentrations above or under 3%, respectively. During the same experiment, one control acquisition in TMN at 30 fps was done before, in the middle and after acquisitions in PVP. For mucin samples, two 1.7-minute recordings were obtained at 15 fps. For growth media tests, two to three recordings of 1 to 3 minutes were

obtained at 30 fps. Numbers of biological and technical replicates, as well as duration of acquisitions, are summarized in Supplementary Table 1.

### Data analysis

Video recordings were binned by a factor of 2x2 by averaging counts and then subjected to a background correction procedure based on dividing the image by a pixel-wise median computed across a sliding window of 101 frames, except for data acquired for mucin experiments, where a sliding window of 41 frames was used. 3D Trajectories were extracted from phase contrast recordings using a high-throughput 3D tracking method based on image similarity between bacteria and a reference library(20). 3D bacterial trajectories were extracted in a tracking volume of approximately 350 µm x 300 µm laterally (x, y) and 200 µm in depth (z) for typically several dozen individuals at a time. Positions were smoothed using 2^nd^ order ADMM-based trend-filtering(70) with regularization parameter *λ* = 0.3 unless stated otherwise (see Supplementary Table 1), and three-dimensional velocities computed as forward differences in positions divided by the time interval between frames. All trajectories with an average speed below a 20 µm/s threshold, unless stated otherwise, were deemed non-motile and discarded. For PVP polymer solutions, the non-motile threshold was adjusted to the population’s swimming speed (see Supplementary Table 1). For samples with 1.3 and 7.4 mM Na^+^, we used *λ* = 0.8 and a non-motile threshold of 10 µm/s.

### Run-reverse-flick analysis

The bacteria’s motility behavior in M9MM was analyzed based on trajectories with a duration of at least 1 s, totaling 23,062 trajectories (58,029 s) and 8,017 trajectories (19,113 s) for strain O395-NT and O395, respectively. The turning event detection is based on the local rate of angular change, computed from the dot product between the sums of the two consecutive velocity vectors preceding and subsequent to a time point. The threshold for a turn to begin is an α-fold rate relative to the median rate of angular change during the trajectory’s run segments, as determined in three iterations of the procedure. We determined by visual inspection of trajectories that a factor α = 8 gave satisfactory results. A turn ends when the local rate of angular change is below the threshold again. The 3D turning angle for a turn beginning at frame i and ending at point j is computed as the angle between the sum of the instantaneous velocity vectors at frames i-2 and i-1 and the sum of those at frames j and j+1. Turning events were labeled flicks or reversals if the turning angle was below 140° or above 150°, respectively. The bacterial orientation during runs (forward/backward) was assigned based on the identity of the two bordering turning events. Backward and forward runs were identified as runs with a flick respectively at the end or at the beginning of the run and a reversal at the other end of the run. For the motility analysis of the O395-NT strain, a total of 8,180 backward and 4,581 forward runs were identified out of 18,533 runs, within a subpopulation of 5,932 trajectories (20,299 s cumulative duration). For the motility analysis of the wildtype O395 strain, 2,283 backward and 1,240 forward runs were identified out of 5,425 runs, within a subpopulation of 1,765 trajectories with a cumulative duration of 5,816 s. For experiments on O395-NT in sodium concentrations of 136 mM, 7.4 mM and 1.8 mM, displayed in Figure 2, we detected 2,769, 6,093 and 10,035 turning events with measurable turning angles, respectively, in 2,683, 5,089 and 4,083 trajectories. For O395 in mucin solutions, we obtained and analyzed 2,327 trajectories with a total duration of 6,027 s, containing 2,587 turning events with measurable turning angle, and enabling us to identify 707 backward and 385 forward runs.

### Deceleration analysis

Runs with a duration of more than 0.33 s and an average speed above 30 µm/s were screened for segments of decreased speed, here called “decelerations”. A deceleration begins when the instantaneous speed drops below a threshold for two consecutive frames and ends when the speed is above the threshold again for two consecutive frames. The threshold is defined as a fraction β of the run’s median speed outside of deceleration events, as determined iteratively in two rounds of the detection procedure. We tested a range of β = 0.3 – 0.9 (Supplementary Figure 4). Figure 4 shows results obtained for β = 0.75.

### Polymer solutions

O395-NT was tracked in solutions of the linear polymer polyvinylpyrrolidone (PVP K90, Sigma 81440, average molecular weight of about 360,000 kDa) with concentrations of ranging from 0.9 to 6.7 % (w/w) in TMN, or of human MUC5B mucin purified from human saliva(71) (a kind gift of K. Ribbeck) at 1.2% (w/w) in M9MM. 3.3 mg lyophilized mucin was dissolved in 275 µl M9MM by shaking at 250 rpm at 4°C for 5 hours. Two hours before the experiment, the solution was slowly pipetted to further homogenize it, and shaken at room temperature at 300 rpm. Macroscopic viscosity measurements for PVP K90 solutions with concentrations in the 0% - 7.2% range were obtained using two falling-ball viscometers (Gilmont, 2-20 cP range and 10-100 cP range) at room temperature (21°C), after calibration using viscosity standards of 11.6 and 48.0 cP (Parangon Scientific, general purpose standards D10 and N26, respectively). The relationship between PVP concentration and viscosity was extracted as a second order polynomial fit with a forced intercept of 0.98 cP at 0% PVP (Supplementary Figure 5).

### Low sodium motility experiment

We varied the motility medium’s Na^+^ concentration at constant ionic strength by mixing M9MM with a buffer identical to M9MM except that all sodium-containing ingredients were replaced by the equivalent potassium-containing ones. NaCl was replaced by KCl, and Na_2_HPO_4_ was replaced by K_2_HPO_4_ (NoNa-M9MM, see Table 2). The experiments were performed as described for experiments in M9MM, except that the culture was not diluted into the motility medium but pelleted at 2,000 rcf for 8 minutes before gentle resuspension in the appropriate motility medium in order to precisely control the final Na^+^ concentration.

### C. crescentus experiment

*C. crescentus* has a dimorphic life cycle: a stalked cell attached to a surface divides and releases a motile daughter cell. We obtained motile cells by a modified plate-release protocol(72). An overnight culture inoculated from an individual *C. crescentus* (CB15, ATCC 19089) colony, grown on 1.5% agar PYE plates streaked from frozen glycerol stock stored at -80°C, was grown to saturation in 2 ml PYE at 30°C, 200 rpm. The overnight culture was diluted 1:100 in fresh PYE, then grown in volumes of 0.5 ml in a 24-well plate incubated at 30°C without agitation for 24 hours. The wells were washed and refilled with fresh PYE, placed at room temperature on a shaker at 130 rpm for another 24 hours. At that stage, the bottoms of the wells were covered with a dense carpet of stalked cells, continuously producing motile cells. For experiments, a well was rinsed three times with PYE, then 0.5 ml of fresh PYE were placed in the well and removed after 5 minutes, containing the newly separated motile cells. Such cell suspensions were immediately injected into a sample chamber and placed on the microscope. Recordings with a duration of 1.5 min were obtained as for *V. cholerae*. Data analysis was performed as for *V. cholerae*, except a threshold factor α = 6 was used for turn event detection as it seemed to produce more accurate results based on visual inspection.

## Acknowledgments

Purified mucin was a kind gift from Katharina Ribbeck (MIT). *V. cholerae* strains O395 and O395-NT were a kind gift from Edward Ryan (MGH/Harvard Medical School). We thank Chloe M. Wu for advice on mucin experiments. This research was supported by the Rowland Institute as well as the Rowland Institute Postdoctoral Fellow and Undergraduate Program, funded by the Rowland Foundation.

## Data availability

All data and computer codes are available from the authors upon reasonable request.

## Author contributions

K.M.T. and M.G. conceived the research. M.M., A.M., and M.G. performed growth condition tests. A.M. and M.G. performed all other experiments on *V. cholera*. M.G. performed experiments on *C. crescentus* as well as viscosity measurements. M.G. developed the turning event analysis. A.M. and M.G. contributed to performing and evaluating turning event analysis. A.M. noticed deceleration events, M.G. developed and performed deceleration event analysis. M.G. and K.M.T. wrote the manuscript. All authors commented on the manuscript.

